# The lysine-rich intracellular loop and cell type-specific co-factors are required for IFITM3 antiviral immunity in hematopoietic stem cells

**DOI:** 10.1101/2021.04.06.438585

**Authors:** G. Unali, A.M.S Giordano, I. Cuccovillo, Abou Alezz M., L. Apolonia, I. Merelli, M. H. Malim, C. Petrillo, A. Kajaste-Rudnitski

## Abstract

The interferon-induced transmembrane protein 3 (IFITM3) inhibits lentiviral gene therapy vector entry into hematopoietic stem cells and can be overcome by Cyclosporine H (CsH), but underlying mechanisms remain unclear. Here, we show that mutating the evolutionarily conserved lysines of the IFITM3 intracellular loop abolishes its antiviral activity without affecting either its localization or its degradation by CsH through non-canonical lysosomal pathways. When confined to the plasma membrane, the lysine-competent IFITM3 lost restriction against VSV-G pseudotyped viral vectors but gained antiviral activity against vectors that fuse directly at the plasma membrane. Interestingly, altering the lysines did not alter IFITM3 homodimerization but impacted higher-order protein complex formation, suggesting loss of interaction with cellular co-factors. In agreement, IFITM3 expression was not sufficient to restrict viral vectors in myeloid K562 cells as opposed to promonocytic THP1 or primary HSC. We exclude the involvement of previously identified factors affecting IFITM3 biology and propose a novel model for IFITM3 restriction that depends on the presence of cellular co-factor(s) that may interact with IFITM3 through the intracellular loop lysine residues. Overall, our work provides significant insight into the mechanisms of action of IFITM3 and CsH that can be exploited for improved gene therapies and broadly acting antiviral strategies.

## Introduction

The low gene manipulation efficiency of human hematopoietic stem and progenitor cells (HSPC) remains a major hurdle for sustainable and broad clinical application of innovative therapies for a wide range of disorders. This remains a great challenge for the field as high vector doses and prolonged *ex-vivo* culture conditions are still required to reach significant transduction levels in clinically relevant human HSPC, imposing costly large-scale vector production and potentially compromising their preservation in culture.

Antiviral innate immunity may partly contribute to HSPC resistance to gene transfer (1). HSPC can directly sense and respond to viral infections (2–4). However, the mechanisms governing these responses have been only partly addressed to date. Interestingly, somatic stem cells ubiquitously express an array of interferon-stimulated genes (ISG) for intrinsic protection from pathogens (5). In agreement, we have recently shown that the interferon-induced transmembrane protein 3 (IFITM3) constitutively blocks VSV-G mediated lentiviral vector (LV) entry into HSPC and can be efficiently counteracted by the non-immunosuppressive Cyclosporine H (CsH) (6).

Together with IFITM1 and 2, IFITM3 is a member of the interferon-inducible transmembrane proteins family of small proteins that are involved in the inhibition of a wide spectrum of viruses (7, 8). Initial studies showed that IFITM3 silencing leads to increased susceptibility to influenza A virus (IAV) in mammalian cells and its over-expression could also impair infection by other influenza virus strains as well as other viruses including the current pandemic-causing SARS-CoV2 (9–15). IFITM3 is a small protein of 144 amino acid. Many topological studies suggest that IFITM3 is a type IV membrane protein, consisting of five domains: a N-terminal domain, an intramembrane domain (IMD), a conserved intracellular loop (CIL), a transmembrane domain (TMD), and a C terminus domain facing the outside of the cell (16–18). IMD and CIL comprise the most evolutionarily conserved domain, known as CD225, in which amino acidic modifications have been associated with altered IFITM3 restriction activity (15, 19).The identification of several post-translational modifications (S-fatty-acylation of Cys71, Cys72 and Cys105, phosphorylation of Tyr20, ubiquitination of Lys24, Lys83, Lys88 and Lys104 and methylation of Lys88) involved in the regulation of IFITM3 antiviral mechanisms further support this IFITM3 topology (18). In mammalian cells, IFITM2 and IFITM3 are mostly present in the early and late endocytic vesicles and on lysosomes (18, 20, 21) whereas IFITM1 lacks the endo-lysosomal sorting motif (YXXΦ) present in both IFITM2 and IFITM3 and is mainly confined at the plasma membrane (7, 12). The cellular localization of the IFITMs may contribute to their viral specificity, as IFITM1 mainly blocks viruses that enter through plasma membrane fusion, while IFITM2 and IFITM3 preferentially inhibit viruses entering via the endocytic route.

Although our understanding of IFITM3 antiviral activity has greatly improved, there is still no consensus regarding its precise molecular mechanism of action and this may differ depending on the target cell and virus. IFITMs inhibit infection at early stages, impeding viral entry into the host cells (7, 22). Viral entry is thought to be impaired via IFITM-mediated changes in the physical properties of the host cell membranes that include increasing curvature, decreasing fluidity and altering membrane composition, thus inhibiting virus-cell membrane fusion (14, 23–25). This restriction seems also to be amplified by incorporation of IFITM proteins into viral particles (23, 26, 27). IFITM1–3 expression has also been shown to consistently reduce viral particle production from cells (8, 23). In addition, IFITMs have been described to suppress HIV-1 protein synthesis by impeding viral mRNA transcription in the polysomes (28).

Here, we have investigated the molecular determinants of IFITM3-mediated inhibition of viral gene therapy vectors in human hematopoietic cell lines as well as in primary HSPC and addressed how CsH induces its degradation. Our work uncovers the requirement of co-factors and a critical region for IFITM3 antiviral activity and provides insight into the proteolytic pathways involved in CsH-mediated counteraction of this antiviral protein.

## Results

### CsH degrades IFITM3 through lysosomal pathways

We have previously shown that CsH increases LV transduction through transient degradation of IFITM3 in HSPC but the mechanisms remain unclear (6). As Cyclosporins are known to bind cyclophilins (Cyp) (Quesniaux et al, 1987), we firstly assessed whether knock-out of the known targets Cyp A, D or F could affect IFITM3 restriction or CsH-mediated inhibition of IFITM3 in THP1 (**Fig. 1A-B**). However, knock-out of Cyclophilins did not affect LV transduction sensitivity to IFNα or CsH (**Fig. 1A-B**). These data suggest that IFITM3 and CsH-mediated effects in THP1 are independent from Cyclophilins.

**Figure 1.**
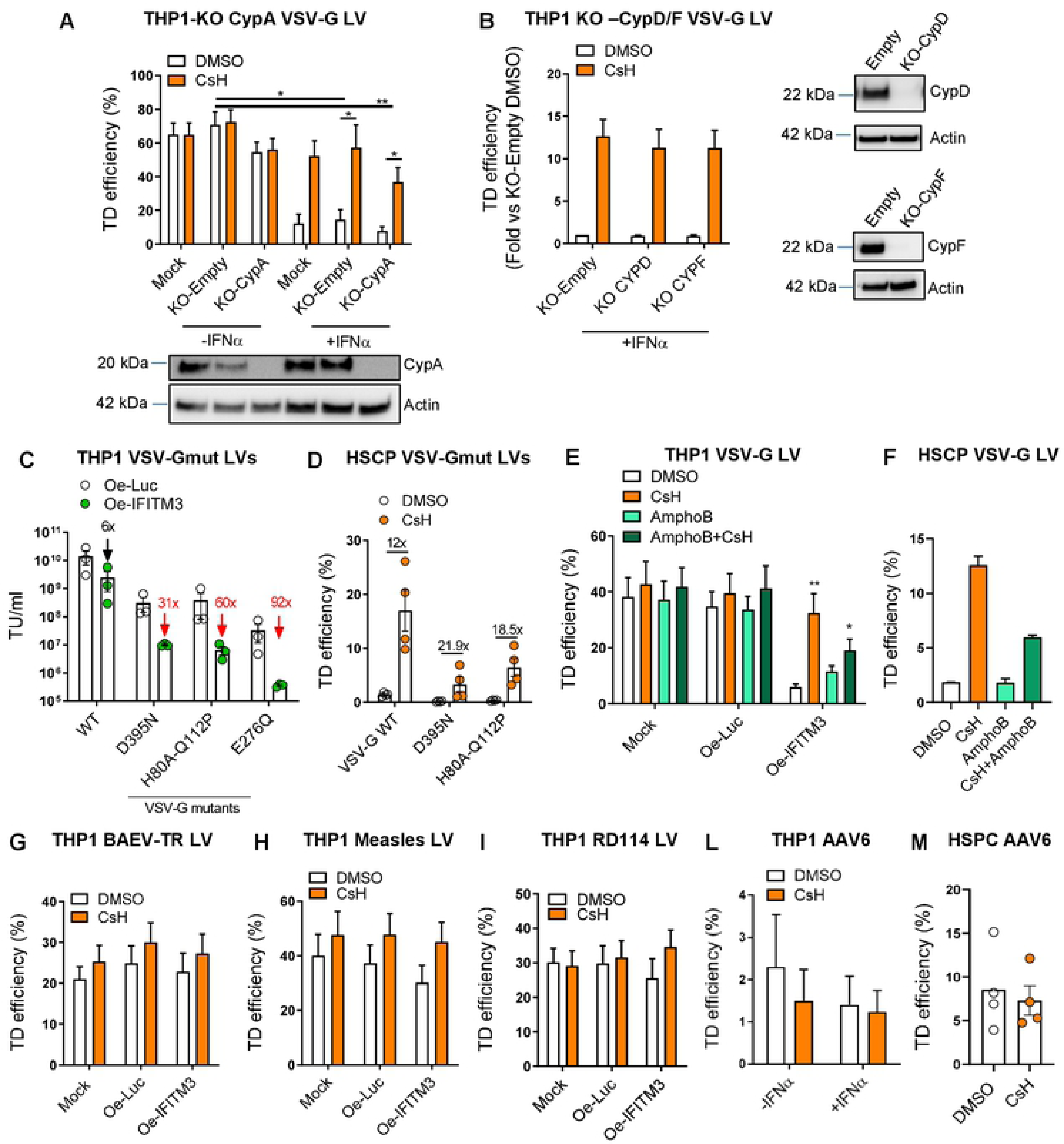
IFITM3 inhibits the endo-lysosomal dependent entry of VSV-G independently of the membrane destabilizing Amphotericin B. LV A-B. THP1 were transduced with a KO-LV expressing both Crispr-Cas9 protein and RNA guide against Cyclophilin A (A), Cyclophilin D or F (B) and then re-challenged with a vector after 24h of stimulation with IFNα. Values are presented as mean ± SEM (n=5-3), p values are for Mann Whitney test. * for p<0.05, **for p<0.01. Cyclophilins knock-out was verified by WB at the time of transduction (TD) (n=3). C. VSV-G wild-type (WT) and mutant LV titers were measured by FACS analysis in THP1 over-expressing IFITM3 and controls. Data are presented as mean ± SEM (n=3). D. CB-HSPC were transduced with VSV-G wild-type or mutant LV in presence or absence of CsH. Transduction efficiencies were measured by FACS five days after TD. Data are presented as mean ± SEM (n=3). E-F. THP1 over-expressing IFITM3 or controls (E) and HSPC were transduced with VSV-G LV in presence or absence of CsH, Amphotericin B or the combination of the two compounds. TD levels were evaluated at FACS 5 days post TD. Data are shown as mean ± SEM (n=5-2), p values are for Mann Whitney test. * for p<0.05, **for p<0.01. G-M. THP1 over-expressing (Oe) IFITM3 and controls were transduced with BaEV-TR (n=9) (G), Measles (n=7) (H) or RD114 (n=6) (I) envelope-pseudotyped LV at MOI 1 and AAV6 (n=5) (L) at MOI 10000. Transduction levels were assessed by FACS five days after TD for LV and three days after TD for AAV6. M. AAV6 transduction was evaluated in CB-HSPC treated or not with CsH three days post-TD (n=4). Data are presented as mean ± SEM (n=4).

IFITM3 has been reported to impede viral egress from the endo-lysosomal compartments (12). In agreement, we observed that LV pseudotyped with mutant VSV-G glycoproteins that are less efficient in fusing with the endo-lysosomal membranes were severely impaired in THP1 over-expressing IFITM3 and HSPC (**Fig. 1C-D**). Moreover, these vectors were the ones to benefit the most from CsH that restored their transduction to the levels of those of a LV pseudotyped with wild-type VSV-G envelope. However, altering lipid membrane stability through Amphotericin B did not rescue IFITM3 restriction (**Fig.1E**) as opposed to previous observations for IAV in the lung epithelial A549 cell line (29, 30), suggesting distinct mechanisms of action in this context. In addition, IFITM3 did not inhibit vectors pseudotyped with glycoproteins fusing directly at the plasma membrane such as BaEV-TR, RD114 or Measles envelopes that remained also insensitive to CsH-mediated enhancement of transduction (**Fig. 1E-F-G**). Of note, we also confirmed that the non-enveloped vector AAV6 remains insensitive to IFITM3 restriction and CsH in HSPC despite entering through endocytosis (**Fig. 1H-I**) (6).

In order to dissect where CsH-mediated degradation of IFITM3 takes place in the cell, we blocked the two best-characterized pathways involved in protein degradation, namely the proteasomal and lysosomal degradation (31). Neither increased transduction nor IFITM3 degradation were affected by addition of the proteasome inhibitor MG132 in THP1 or HSPC (**Fig. 2A-C; 2H**). Moreover, in agreement with previous reports (32), proteasomal inhibition enhanced LV transduction in HSPC and the effect was additive to that of CsH (**Fig. 2B**), likely indicating that MG132 may act on a different restriction block that we predict is independent from IFITM3. In contrast, inhibition of lysosomal degradation by Bafilomycin was sufficient to block both CsH enhancement of transduction as well as IFITM3 degradation (**Fig. 2D-F; 2H-I**). Consistent with a block of IFITM3 turnover in the endo-lysosomes we observed a significant increase in IFITM3 protein expression in THP1 and HSPC treated with Bafilomycin (**Fig. 2G-I**). This effect, in combination with the capacity of Bafilomycin to inhibit acidification of endo-lysosomes, may explain the strong inhibition of LV transduction observed after Bafilomycin exposure (**Fig. 2D-E**). Because Bafilomycin inhibits also early and late autophagy we tested whether autophagy inhibition through different compounds may lead to the same observations. However, neither 3MA nor Spautin-1 treatment altered CsH-mediated rescue of transduction in THP1 or HSPC (**Fig. S1 A-D**), indicating that lysosomal-specific degradation pathways are involved in the CsH-mediated degradation of IFITM3.

**Figure 2.**
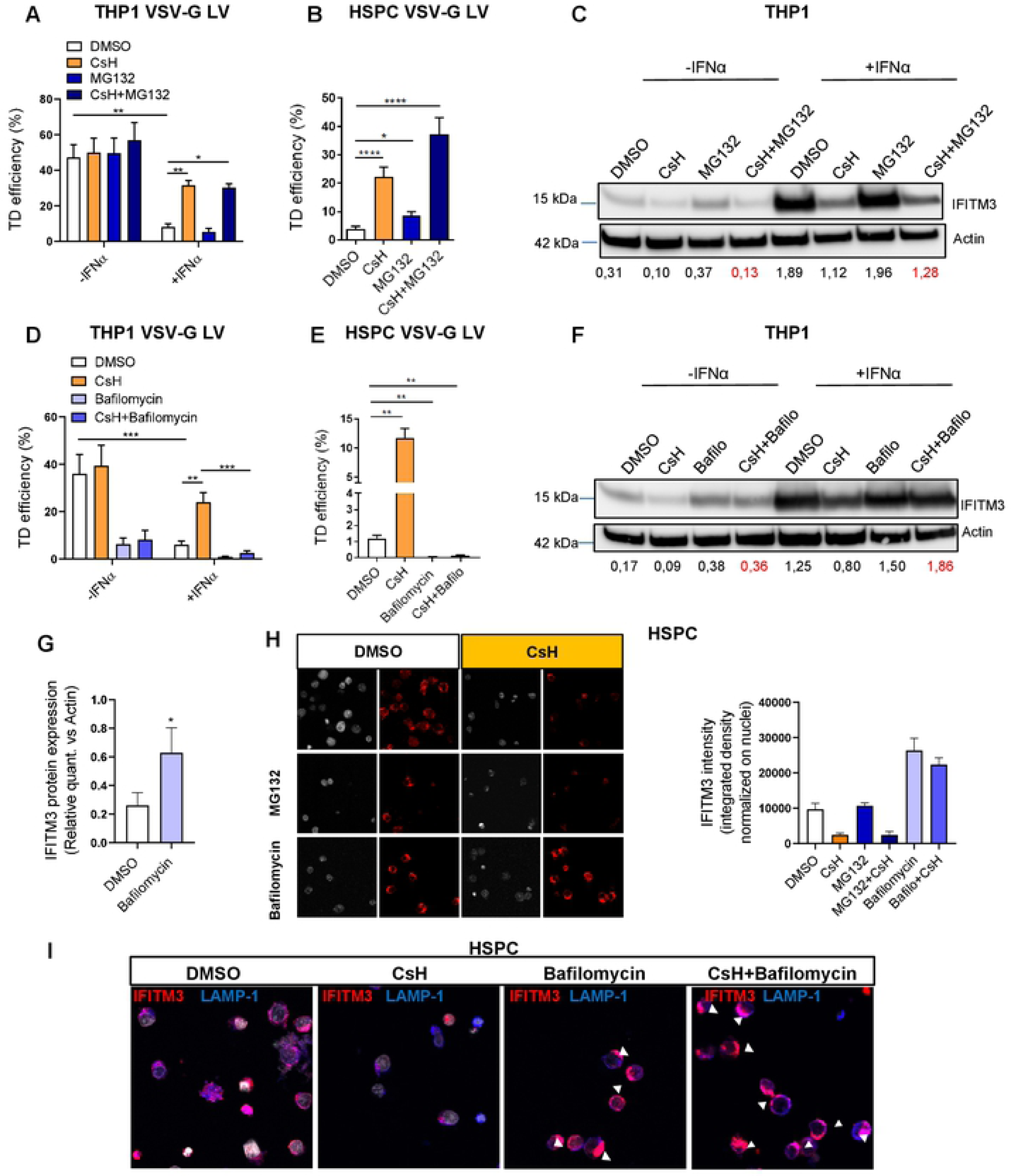
CsH targets IFITM3 for lysosomal degradation. **A-B-D-E**. THP1 pre-stimulated or not for 24h with IFNα (**A-D**) and human CB-HSPC (**B-E**) were transduced with VSV-G LV in presence or absence of CsH, the proteasomal inhibitor MG132, the lysosomal inhibitor Bafilomycin or the combination of CsH with one of the two inhibitors. Transduction efficiency was evaluated at FACS five days post-TD. Data are presented as mean ± SEM (n=6-3), p values are for Mann Whitney test,* for p<0.05, **for p<0.01, ***for p<0.001, ****for p<0.0001. **C-F.** IFITM3 protein expression was measured by WB after an O/N treatment with CsH, MG132, Bafilomycin or the combination of CsH with one of the two inhibitors in THP1 pre-stimulated with IFNα, one representative image is shown. IFITM3 was quantified and normalized to Actin using ImageJ (n=3-5). **G.** IFITM3 protein was quantified in THP1 treated or not with Bafilomycin using ImageJ, normalization was performed over Actin. Data are shown as mean ± SEM (n=6), p values are for Wilcoxon test, * for p<0.05. **H.** Immunofluorescence images were performed using TCS SP5 Leica confocal microscope, 60x with oil on human HSPC treated or not with CsH, MG132, Bafilomycin or the combination of CsH with one of the two inhibitors. Representative zoomed images are shown (n=6). IFITM3 integrated density was quantified by ImageJ. **I.** Co-localization of IFITM3 (in red) with the lysosome associated membrane protein 1 (LAMP1) marker (in violet) was evaluated in HSPC by immunofluorescence using TCS SP5 Leica confocal microscope 60x with oil (n=6 images).

Noteworthy, we observed that the plasma-membrane confined IFITM3 Y20 phosphomutants that we have previously shown to have lost antiviral activity against VSV-G LV in HSPC (6) inhibited plasma-membrane (PM) fusing vectors such as BaEV, RD114 and Measles pseudotyped LV (**Fig. 3A**). However, the Y20 mutant was resistant to CsH degradation (**Fig. 3B**), which we suggest explains why CsH could not rescue its antiviral activity against PM-fusing LV in THP1 (**Fig. 3A**). The Y20 mutant insensitivity to CsH could be ascribed to its altered cellular localization (**Fig. 3B**) as appropriate trafficking is required for protein degradation. In this regard, Y20 phosphorylation by the Fyn kinase has been shown to regulate IFITM3 trafficking in HEK293 cells (33). However, VSV-G LV were still susceptible to IFNα antiviral activity in THP1 KO for Fyn and the PP2 inhibitor of Src-kinases did not affect IFITM3-mediated inhibition of LV transduction in THP1 and HSPC (**Fig. 3C-E**), indicating that phosphorylation by Fyn is not required for IFITM3 restriction in our experimental settings.

**Figure 3.**
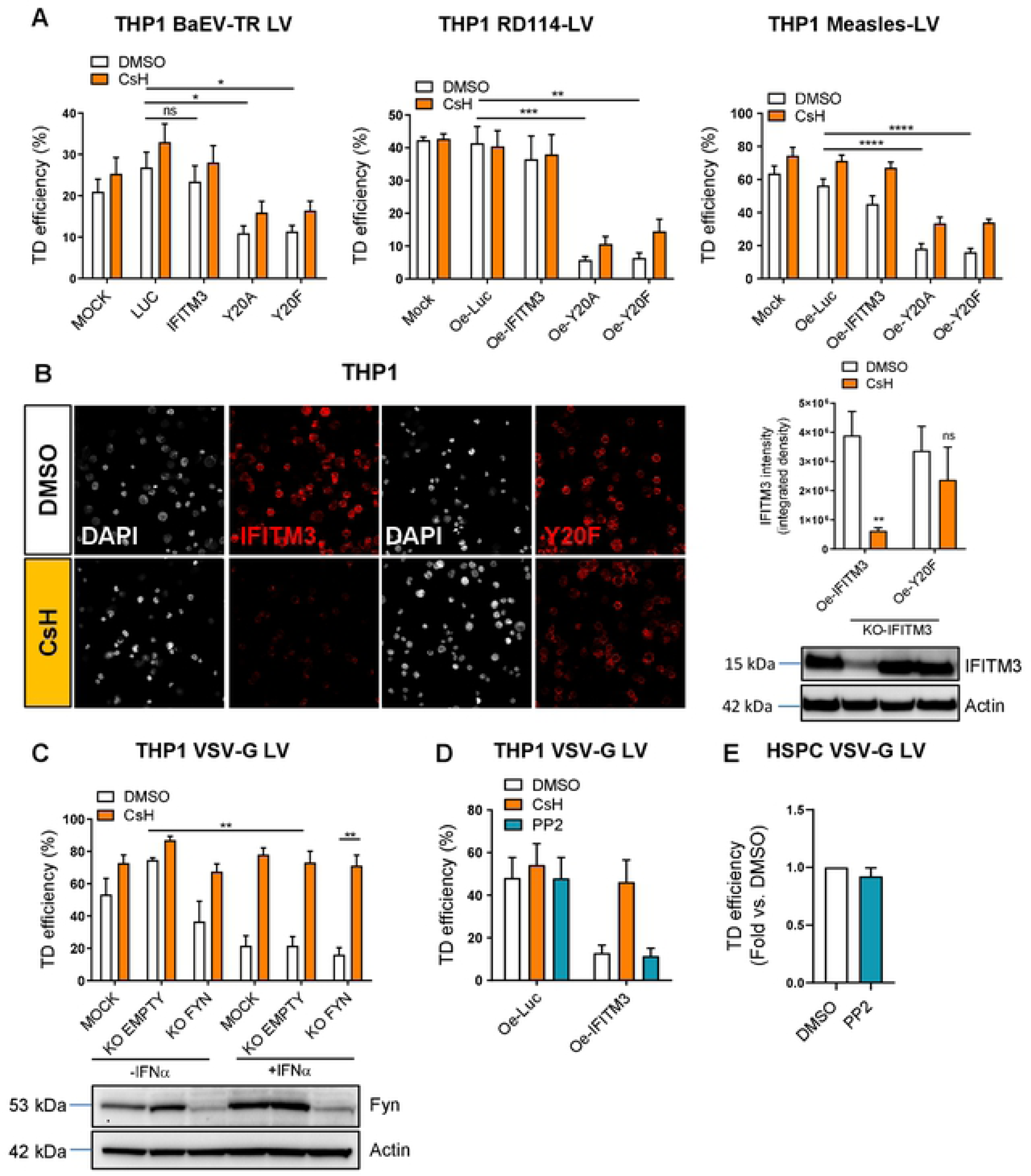
Altered IFITM3 localization redirects its antiviral activity and impacts its sensitivity to CsH. **A.** THP1 over-expressing IFITM3 wild-type, IFITM3-Y20 mutants and controls were transduced with BaEV-TR LV, RD114 LV and Measles LV at MOI 0,5-1 with or without CsH. Transduction efficiency was measured by FACS five days after transduction. Data are presented as mean ± SEM (n=6-5), p values are for Mann Whitney test, * for p<0.05, **for p<0.01, ***for p<0.001, ****for p<0.0001. **B.** Immunofluorescence images were performed using TCS SP5 Leica confocal microscope, 60x with oil on THP1 over-expressing IFITM3 wild-type or IFITM3 phosphomutant Y20F after 16h of CsH treatment. IFITM3 integrated density was quantified by ImageJ. Zoomed images are shown. Data represent the mean ± SEM of four independent experiments (n=18 images), p values are for Mann Whitney test. **for p<0.01, ****for p<0.0001. IFITM3 degradation was assessed also by WB (n=4). **C.** THP1 Knock-out for Fyn and controls were pre-stimulated with IFNα for 24h and then transduced with VSV-G LV in presence or absence of CsH. Transduction was measured by FACS five days later. Data are shown as mean ± SEM (n=4), p values are for Mann Whitney test, **for p<0.01. **D-E.** THP1 over-expressing IFITM3 and controls **(D)** and HSPC **(E)** were transduced with VSV-G LV in presence or absence of CsH or the Src-kinase inhibitor PP2. Transduction was evaluated at FACS after five days. Data represent the mean ± SEM (n=5-3).

### CsH-mediated degradation of IFITM3 occurs independently from single lysine ubiquitination and methylation

The NEDD4 E3-ubiquitin ligase has been described as responsible for IFITM3 degradation and turn-over by specific ubiquitination of the lysine in position 24 (34, 35). To address whether CsH could cause IFITM3 degradation through NEDD4-mediated ubiquitination we tested its effect on an IFITM3 mutant (ΔProl) devoid of the domain required for interaction with NEDD4 (34) that we over-expressed in THP1 devoid of endogenous IFITM3. The ΔProl IFITM3 retained CsH susceptibility (**Fig. 4A-B**), demonstrating that this domain is not required for CsH-mediated degradation of IFITM3. In addition, compared to the wild-type, this mutant has been reported to harbor exacerbated antiviral activity, explained by an extended half-life and a consequent accumulation in the endo-lysosomes (34, 36). However, we observed that both wild-type and mutant IFITM3 shared similar antiviral activity and comparable localization in the endo-lysosomal compartments in (**Fig. 4C**).

**Figure 4.**
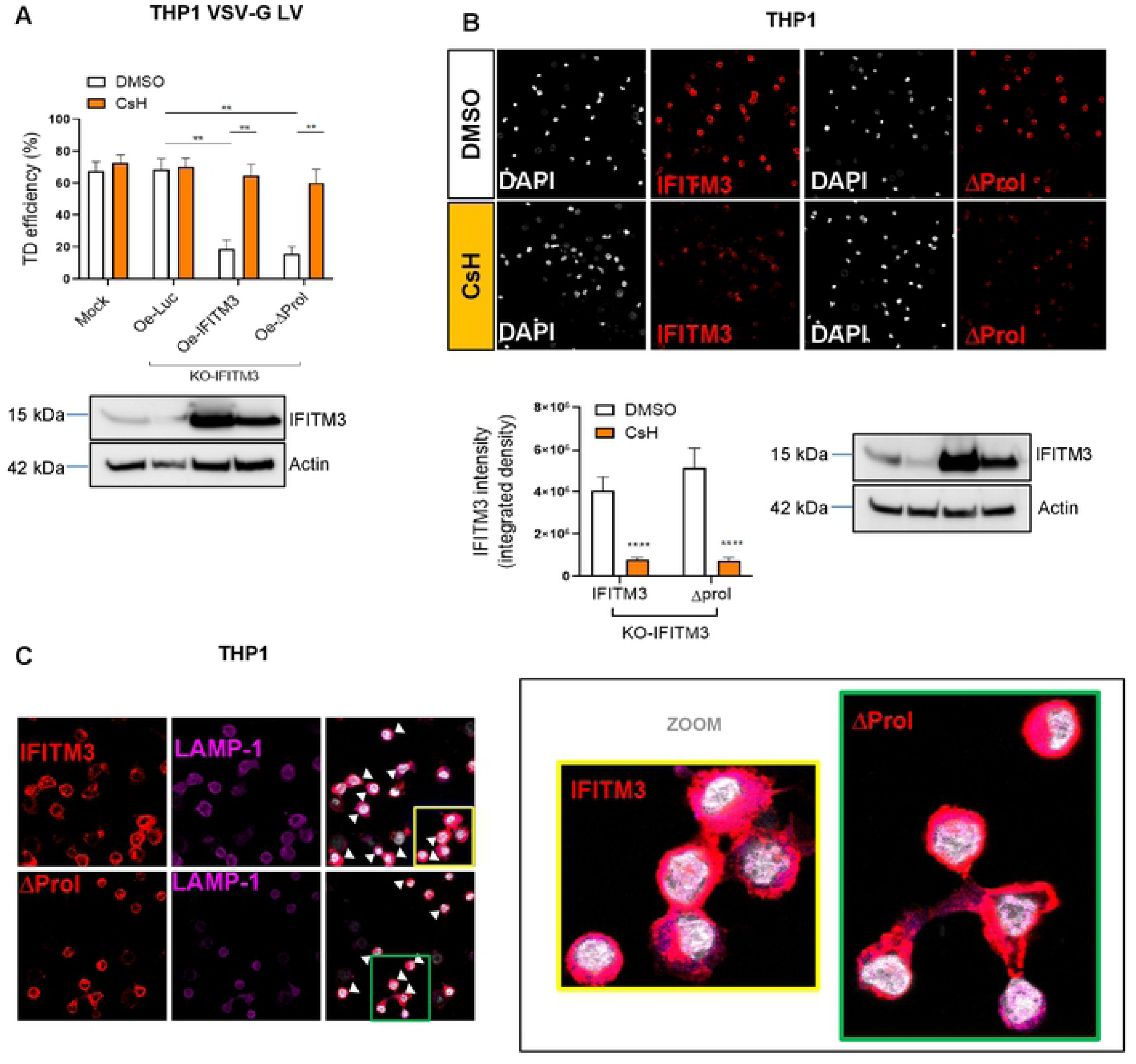
IFITM3 devoid of the NEDD4 interaction domain remains susceptible to CsH. **A.** THP1 over-expressing (Oe) IFITM3 wild-type, IFITM3 defective for the interaction domain with the ubiquitin ligase NEDD4 (ΔProl) or control were transduced at MOI 1 with VSV-G LV. Transduction efficiency was evaluated by FACS five days post-TD. Data represent the mean ± SEM (n=7), p values are for Mann Whitney test **for p<0.01. IFITM3 protein expression was measured by WB at time of transduction (n=4). **B.** IFITM3 wild-type and IFITM3 ΔProl degradation by CsH was assessed by IF and WB in THP1 after 16h of CsH treatment. IFITM3 was quantified through ImageJ. Data represent the mean ± SEM of four independent experiments (n=12 images), p values are for Mann Whitney test. ****for p<0.0001. **C.** Co-localization (purple areas marked by white arrows) of IFITM3 wild-type and ΔProl proteins (in red) with the lysosome associated membrane protein 1 (LAMP1) marker (in violet) was evaluated by immunofluorescence using TCS SP5 Leica confocal microscope 60x with oil (n=12 images). Zooms of the yellow and green boxed areas are shown on the left.

Ubiquitination of the lysine residues is central for protein degradation (37). In agreement, IFITM3 is reported to be degraded predominantly in response to ubiquitination of lysine 24 and to a minor extent of the other lysines in position 83, 88 and 104 (18, 21, 34–36, 38). To understand if CsH could preferentially require one of the lysine residues to degrade IFITM3 we generated single lysine to arginine mutants that we over-expressed in THP1. All mutants were capable of inhibiting VSV-G LV transduction (**Fig. 5A**). Accordingly, no major differences in endo-lysosomal localization were observed (**Fig. 5B-S2**). Importantly, all mutants were also degraded by CsH (**Fig. 5C**). Moreover, these results likely exclude the involvement of methylation in CsH-mediated degradation of IFITM3 as the lysine at position 88 has been recently shown to be a target of methylation by Set7, exploited by both VSV and IAV to down-regulate IFITM3 protein (8, 39). Moreover, IFITM3 88K/R did not show improved capacity of inhibiting VSV-G LV (**Fig. 5A**), suggesting that lysine 88 methylation does not play a major role in inhibiting IFITM3 antiviral activity against VSV-G LV transduction in THP1 cells.

**Figure 5.**
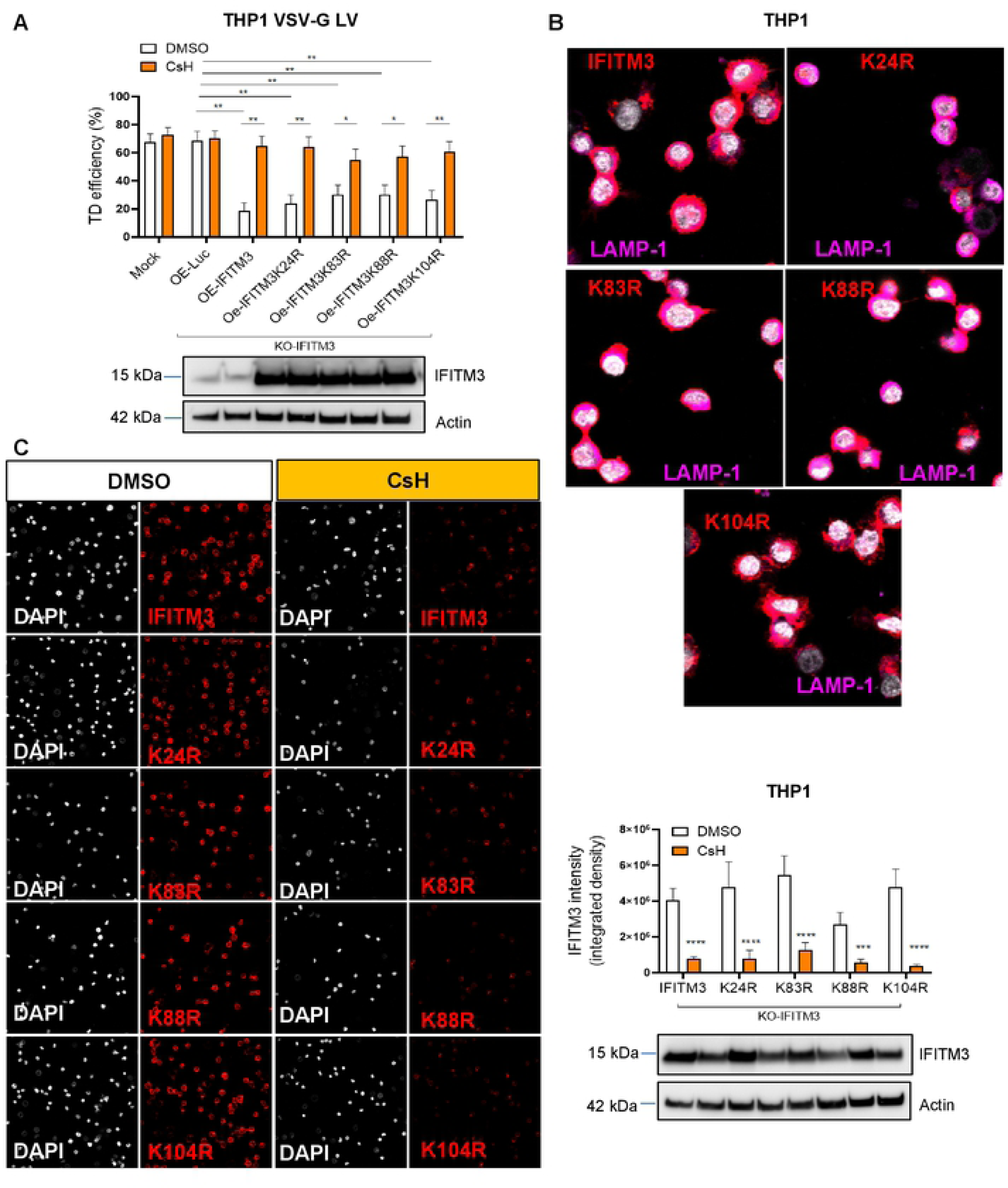
CsH-mediated degradation of IFITM3 is independent from single lysine ubiquitination. **A.** THP1 over-expressing (Oe) IFITM3 wild-type, IFITM3 lysine to arginine mutants or control were transduced at MOI 1 with VSV-G LV. Transduction level was measured at FACS five days after transduction. Data represent the mean ± SEM (n=7), p values are for Mann Whitney test. * for p<0.05, **for p<0.01. IFITM3 protein levels were assessed by WB at time of transduction. **B.** IFITM3 wild-type and single lysine mutants co-localization with LAMP-1 was evaluated by IF. Immunofluorescence images were acquired using TCS SP5 Leica confocal microscope, 60x with oil. Zoomed images are shown (n=12). **C.** Immunofluorescence images were performed using TCS SP5 Leica confocal microscope, 60x with oil on THP1 over-expressing IFITM3 or IFITM3 single lysine mutants after 16h treatment with CsH. IFITM3 intensity was quantified by ImageJ. Data represent the mean ± SEM of five experiments (n=12), p values are for Mann Whitney test. * for p<0.05, **for p<0.01, ***for p<0.001, ****for p<0.0001.

As the infected cells can up-regulate the histone demethylase LSD1 to remove Set7 methylation and promote IFITM3 antiviral activity (40) we also tested a LSD1-specific inhibitor and noted that it did not affect IFITM3 antiviral activity in THP1 cells over-expressing IFITM3 or primary HSPC (**Fig. S3 A-B**). In agreement with specific inhibition, LSD1 protein down-regulation by LSD1 inhibitor lead to increased methylation of the histone H3K9 but did not affect IFITM3 protein expression (**Fig. S3 C**). Taken together, these data invoke a mechanism of degradation of IFITM3 by CsH that does not depend on any single lysine of IFITM3 nor Set7-mediated methylation.

### The lysine residues in the CD225 region of IFITM3 and cell-type specific co-factors are required for antiviral activity

To determine whether CsH-mediated degradation of IFITM3 could occur through ubiquitination of any one of the multiple lysines, we generated an IFITM3 mutant lacking all the lysine residues (**Fig. 6A**). Arginines were selected for lysine substitution in order to maintain the basic charges and preserve conformation. Strikingly, and in contrast to previous reports regarding a similar lysine-less mutant (18, 36), our IFITM3Lys/Arg mutant completely lost antiviral activity against VSV-G LV in THP1 cells (**Fig. 6B**). Trafficking alterations of the mutant were excluded by localization studies performed in THP1 or in primary human monocyte-derived macrophages (MDM) (**Fig. 6C-D**) in which CsH-mediated enhancement of transduction can be recapitulated upon type I IFN stimulation (**Fig. 6E**). Remarkably, CsH was still capable of degrading this lysine-less mutant, indicating that an alternative degradation pathway(s) is involved in CsH triggered proteolysis (**Fig. 6F-G, S4**).

**Figure 6.**
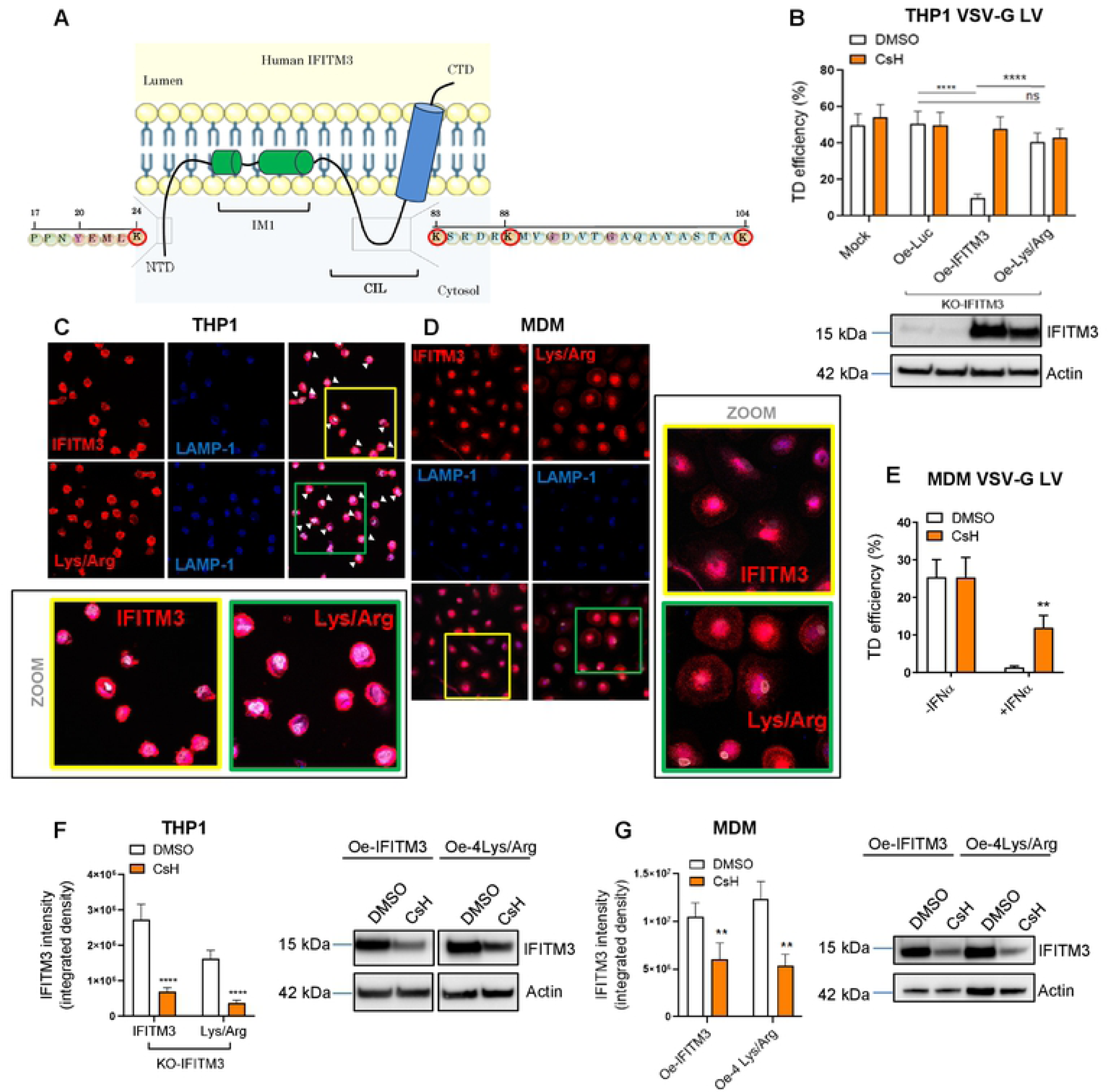
Lysine-less IFITM3 mutant loses antiviral activity but remains susceptible to CsH. **A.** Scheme of IFITM3 protein configuration. The lysine residues mutated into arginines are highlighted in red. **B.** THP1 over-expressing (Oe) IFITM3 wild-type, IFITM3 lysine-less mutant or control were transduced at MOI 1 with VSV-G LV. IFITM3 protein expression was evaluated by WB at time of TD (n=4). Transduction level was measured by FACS. Error bars represents the mean ± SEM (n=8), p value are for Mann Whitney test. ****for p<0.0001. **C-D.** Immunofluorescence images were performed using TCS SP5 Leica confocal microscope, 60x with oil on THP1 **(C)** or MDM **(D)** to assess co-localization (purple area marked by white arrows) of IFITM3 and IFITM3 lysin-less mutant (in red) with the lysosomal marker LAMP1 (in blue). Zoomed images are shown for THP1. **E.** Human monocyte-derived macrophages (MDM) were pre-stimulated with IFNα for 24h and then transduced with LV at MOI 1. Transduction efficiency was measured by FACS five days after TD. Data are shown as mean ± SEM (n=7) **F-G.** IFITM3 or IFITM3 lysine-less degradation by CsH was investigated by IF in THP1 and MDM after 16h treatment with CsH. CsH-mediated degradation of IFITM3 was also confirmed by WB in THP1. IFITM3 protein expression was quantified by ImageJ. Data represent the mean ± SEM of at least three experiments (n>12), p values are for Mann Whitney test. **for p<0.01, ****for p<0.0001.

NMR studies suggest that lysine-less IFITM3 has no topological changes, as the lysines are in an intra-cytosolic portion that does not span the lipid membrane (16). Therefore, we reasoned that ablation of the antiviral phenotype should be independent of changes in the protein conformation. Studying the positioning of the mutated lysines we noticed that Lys83, Lys88 and Lys104 clustered together in the intracellular loop (CIL), known to be part of the most evolutionarily conserved CD225 domain of IFITM3 (19) (**Fig. 6A**). Similarly, we observed that the three lysines are conserved among the antiviral IFITMs and maintained across species, pointing towards an evolutionarily conserved functional role of these amino acids (**Fig. 7A**). Because amino-acidic changes in the CD225 domain have been shown to potentially affect IFITM3 dimerization and consequently antiviral activity (19, 41, 42), we assessed whether the loss of restriction could be caused by defects in IFITM-IFITM interaction. The lysine-less mutant maintained dimerization capacity in non-denaturating electrophoresis, likely excluding a lack of dimerization as the mechanism behind loss of antiviral activity of the lysine-less IFITM3 (**Fig.7B**). However, we observed a difference in higher order protein complex formation with increased ratio of IFITM3 dimers over higher molecular weight complexes (**Fig. 7B**), suggesting potential loss of co-factor interactions necessary for IFITM3 antiviral activity. In agreement, over-expression of IFITM3 was not sufficient to restrict LV transduction in the myeloid cell line K562 (**Fig.7C**). Of note, also IFNα failed to inhibit transduction in these cells despite readily inducing expression of IFN-stimulated genes, including endogenous IFITM3 (**Fig. 7D**). Defects in IFITM3 localization were excluded by IF analysis (**Fig. 7E**). Restoring the lysine in position 24 did not re-establish IFITM3 restriction in THP1 cells (**Fig. 7F**) indicating that the critical region for antiviral restriction is confined to the small intracellular loop of IFITM3. Of note, the triple lysine mutant IFITM3 readily localized within the lysosomes and remained sensitive to CsH-mediated degradation (**Fig. 7F-G, S5A-B**), highlighting that distinct residues and regions of the protein are involved in restriction and turnover. Taken together, our data identify the three lysine residues within the intracellular loop of IFITM3 as critical for its antiviral activity against VSV-G pseudotyped LV but dispensible for its degradation by CsH, and further suggest that cell-type specific co-factors are required for IFITM3 antiviral activity.

**Figure 7.**
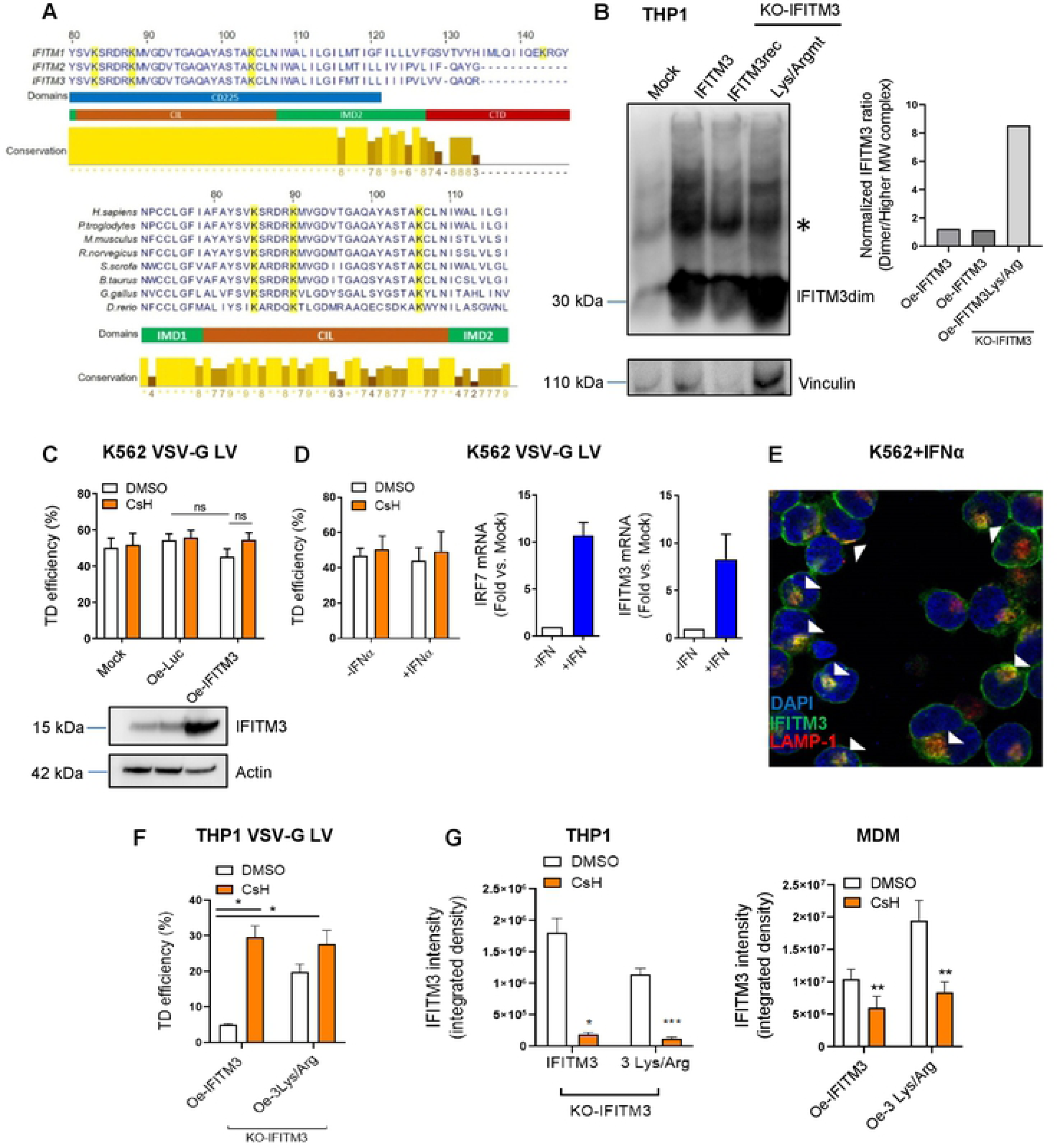
The lysines of the intracellular loop of IFITM3 and cell-type specific co-factors are required for antiviral activity. **A.** Partial protein alignment of the lysine-rich CIL domain of human antiviral IFITMs and IFITM3 across different species**. B.** IFITM3 dimerization and oligomerization was assessed by native PAGE followed by WB analysis in THP1. IFITM3 higher order protein complex bands are highlighted by the asterisk. Normalized ratios between IFITM3 dimers and higher order complexes was quantified by ImageJ. **C.** K562 over-expressing (Oe) IFITM3 or controls were transduced with VSV-G LV (n=7). Transduction levels were assessed by FACS five days after TD. **D.** K562 were pre-stimulated with IFNα for 24h and then transduced with LV at MOI 1. Transduction efficiency was measured by FACS five days after TD. IRF7 and IFITM3 mRNA levels were measured by qPCR to assess ISG induction by IFNα at the time of transduction Data are shown as mean ± SEM (n=3). **E.** IFITM3 localization was analyzed by IF in K562, zoomed images are shown (n=12). **F.** Transduction efficiency was measured by FACS in THP1 over-expressing (Oe) IFITM3 wild-type, IFITM3 triple lysine mutant or control transduced at MOI 1 with VSV-G LV. Data are mean ± SEM (n=5), p values are for Mann Whitney test. * for p<0.05. **G.** Degradation of IFITM3 wild-type and triple lysine mutant IFITM3 was measured after 16h treatment with CsH in THP1 and MDM by IF. Data represent the mean ± SEM of at least three independent experiments (n=12 images), p values are for Mann Whitney test. * for p<0.05, ****for p<0.0001.

## Discussion

Our work sheds significant light on molecular mechanisms of IFITM3-mediated restriction of viral entry and how CsH counteracts it. Among the IFITM family of proteins, IFITM3 has been the best characterized for blocking a broad range of viruses, including HIV-1 (7, 8, 11, 12, 14). Although its antiviral activity is known to be controlled by several post-translational modifications (18), a detailed molecular mechanism of action has not been identified yet. Our data suggests that IFITM3 insensitive vectors fuse at the plasma membrane as opposed to the VSV-G-pseudotyped LV that enters through pH-dependent endocytosis. In agreement, other viruses that share similar entry routes to VSV-G LV have shown susceptibility to IFITM3 restriction (22, 30). Among them, IAV is blocked at the stage of fusion with the endosomal membranes (9, 24, 25, 30). Interestingly, we confirmed that a similar mechanism occurs also upon VSV-G LV transduction in IFITM3 expressing cells, as VSV-G mutants defective for viral fusion with the endo-lysosomal membranes were more susceptible to IFITM3 restriction. However, differently from IAV, the IFITM3 restriction of VSV-G LV could not be rescued by Amphotericin B, suggesting that the mechanism of action does not rely on membrane stability in this context.

In agreement with a block at the level of the VSV-G hemi-fusion with the endo-lysosomes, IFITM3 Y20 phosphomutants that were retained at the plasma-membrane showed loss of antiviral activity against VSV-G LV. Conversely, these mutants gained antiviral activity against LV entering through direct fusion at the cell surface. As these mutants likely resemble IFITM1, known to localize at the plasma-membrane and to restrict viruses fusing therein, IFITM1 could well contribute to inhibiting PM-fusing vectors in HSPC and other restricted primary cells. Of note, IFITM3 did not restrict AAV6 transduction despite the endocytic route used by AAV to enter the cells. Lack of a viral envelope could potentially explain this observation as no hemi-fusion between viral and cellular lipid membranes occurs in this context. Nevertheless, non-enveloped reoviruses that enter through endocytosis have been shown to be susceptible to IFITM3 (43), indicating that more complex mechanisms of selectivity may be involved.

Similarly to mutations of the Y20 residue, phosphorylation by the Fyn kinase in the same tyrosine has been reported to regulate IFITM3 trafficking to the plasma membrane and ubiquitination (18, 21, 35). However, neither IFITM3 antiviral activity nor CsH-mediated rescue of transduction were affected by Fyn depletion in THP1, potentially indicating that other kinases could be involved in regulating IFITM3 restriction of VSV-G LV in these cells. Moreover, we showed that trafficking is necessary for CsH-mediated degradation of IFITM3, as the Y20 mutant IFITM3 was insensitive to CsH. In agreement, CsH could not rescue Y20 IFITM3-mediated antiviral activity against PM fusing-LV transduction. This is in line with the fact that proper trafficking is critical for protein degradation (44, 45). In this context, we have shown that proteasome inhibition by MG132 did not affect CsH-mediated degradation of IFITM3. On the contrary, the combination of MG132 with CsH lead to a synergistic positive effect on transduction in HSPC, indicating that the two drugs have separate mechanisms of action. This data further underscores the potential presence of other antiviral blocks in HSPC in agreement with previous reports of MG132-mediated enhancement of LV transduction in HSPC (32). Interestingly, the immunoproteasome was recently shown to regulate the antiviral activity of TRIM5α (46), a well-known lentiviral restriction factor (47–49). Whether MG132-mediated enhancement of transduction could be related to effects on TRIM5α in HSPC remains to be investigated.

Although we discovered that CsH leads to IFITM3 degradation through lysosomal pathways, independently from autophagy, the mechanism that regulate this process is still under investigation. We excluded involvement of CypA, D and F in CsH and IFITM3 mediated effects as LV transduction was impaired in Cyps KO cells and still benefitted from CsH. Formyl peptide receptor 1 (FPR1) is one of the few characterized targets of CsH (de Paulis et al, 1996). Interestingly, FPR1 is a member of the N-formyl peptide receptors (FPR) family of pattern recognition receptors (PRR) involved in the regulation of innate immune responses (50). Nevertheless, we did not observe major alterations of IFITM3 antiviral activity and CsH–mediated enhancement of LV transduction in cells KO for FPR1 in our previous work (6). We also excluded involvement of NEDD4-dependent or single lysine ubiquitination of IFITM3 and methylation (18, 21, 34, 36, 39, 40) in the CsH-mediated degradation of IFITM3.

Importantly, we show that the lysines in position 83, 88 and 104 located within an intracellular loop of IFITM3 are evolutionarily conserved and necessary for its antiviral activity against VSV-G LV. This is in contrast with previous literature where a similar lysine-less mutant IFITM3 was shown to have increased antiviral activity against IAV due to the loss of its ubiquitin-mediated degradation (36, 51). Possible topological changes caused by these mutations have been excluded in other studies (16). Amino-acidic alterations in neighboring regions of these three IFITM3 lysines have been reported to affect dimerization (19, 42) and antiviral activity. However, our data indicate that the IFITM3 lysine-mutant is still capable of forming dimers, although we observed a lower capacity to form slower migrating IFITM3 protein complexes, including IFITM3 oligomers. It is tempting to speculate that the lysine residues within the intracellular loop of IFITM3 could be an interaction interface for cellular co-factor(s) necessary for its antiviral activities that are expressed in THP1 cells and primary hematopoietic cells such as macrophages and HSPC, but not in K562 cells. It will be of interest to identify these co-factors and explore their role in the virus and cell-type specific effects of IFITM3.

Interestingly, mutations of the lysines in position 83 and 104 have been recently associated with a decreased affinity of IFITM3 for phosphatidylinositol (3,4,5)-3 phosphate, causing loss of amplification of PI3K signals required for B cell activation as well as tumorigenesis (52). We have shown that PI3K inhibitors, such as 3MA, do not affect IFITM3 antiviral activity or its sensitivity to CsH. However, phosphatidylinositols exhibit key roles in endocytic membrane trafficking to and from the plasma membrane and within the endolysosomal compartment (53–55). Therefore, it will be of interest to assess the capacity of the IFITM3 lysine mutant to bind phosphatidylinositol (3,4,5)-3 phosphate (PI-3,4,5-3P) and the contribution of this phospholipid binding to IFITM3 antiviral activity. Remarkably, mutation of all four lysines within IFITM3 did not affect its susceptibility to CsH-mediated degradation, indicating that pathways that act independent of lysine ubiquitination are engaged in the CsH-mediated mechanism of IFITM3 degradation. Cysteine, Serine and Threonine ubiquitination are emerging as alternative protein degradation pathways (56) while the N-degron pathway is an important quality control process to ensure protein degradation (57, 58). Whether such mechanisms could be involved in CsH-mediated degradation of IFITM3 remains to be investigated.

Taken together, our work uncovers novel molecular mechanisms controlling innate immune defenses in human blood cells, including primary hematopoietic stem cells. Understanding the mechanisms of action of IFITM3 and CsH will inform the development of improved gene therapies and of broadly acting antiviral strategies against IFITM–sensitive viruses, including the current pandemic-causing SARS-CoV2.

## Materials and Methods

### Vectors

Third generation LV stocks were prepared, concentrated and titered as previously described (6, 59, 60). Briefly, Self-inactivating (SIN) LV vectors were produced using the transfer vector pCCLsin.cPPT.hPGK.eGFP.Wpre, the packaging plasmid pMDLg/pRRE, Rev-expressing pCMV-Rev and the VSV-g envelop-encoding pMD2.VSV-G plasmids. VSV-G mutants, D395N, H80A-Q112P, E276Q have been previously described (61, 62)and were used in place of the VSV-G envelope-encoding pMD2.VSV-G plasmid. For pseudotyping LV with the mutant baboon retrovirus envelope, the endogenous feline viral envelope RD114 and measles envelopes, pMD2.VSV-G was replaced by the BaEV-TR, RD114 and Measles env encoding plasmids during vector production as previously described (63). Vpx-containing lentiviral vector stocks were produced as previously described (64). AAV6 donor templates for HDR were generated from a construct containing AAV2 inverted terminal repeats, produced by triple-transfection method and purified by ultracentrifugation on a cesium chloride gradient as previously described (65).The bidirectional LV (66)were used to over-express the coding sequence (CDS) of candidate human genes under the control of the human phosphoglycerate kinase (PGK) promoter and the eGFP and the minimal cytomegalovirus (mCMV) promoter forming the antisense expression unit. Bidirectional transfer plasmid for over-expression of IFITM3was made by cloning IFITM3 cDNA sequence into BamHI and XmaI sites. IFITM3 mutants were made either by site-directed mutagenesis PCR or designed and then synthesized by Twin-Helix. KO experiments were performed using lentiCRISPR v2 which encode both Cas9 and RNA guides against the gene of interest as previously described (6). lentiCRISPR v2 against human CypD and CypF were kindly provided by Luis Apolonia.

### Cells

#### Cell lines

The human embryonic kidney 293T cells (HEK293T) were maintained in Iscove’s modified Dulbecco’s medium (IMDM; Sigma), human THP-1 cells were maintained in Roswell Park Memorial Institute medium (RPMI; Lonza). All medium were supplemented with 2-5 or 10% fetal bovine serum (FBS; Gibco) according to the experimental setting, penicillin (100 IU/ml), streptomycin (100 μg/ml) and 2% glutamine.

#### Primary cells

Human CD34^+^ HSPC and CD14^+^ monocytes were isolated through positive magnetic bead selection according to manufacturer’s instructions (Milteney) from umbilical cord blood or from mononuclear cells collected upon informed consent from healthy volunteers according to the Institutional Ethical Committee approved protocol (TIGET01). Otherwise, CB, bone marrow (BM) or G-CSF mPB-CD34^+^ cells were directly purchased from Lonza or Hemacare. MDM were differentiated from isolated CD14^+^ monocytes in Dulbecco’s modified eagle’s medium (DMEM) supplemented with 10% FBS, penicillin (100 IU/ml), streptomycin (100 μg/ml), 2% glutamine and 5% human serum AB (Lonza) for seven days. All cells were maintained in a 5% CO_2_ humidified atmosphere at 37°C.

#### Transduction

Human CB-derived HSPC were cultured in serum-free StemSpan medium (StemCell Technologies) supplemented with penicillin (100 IU/ml), streptomycin (100 μg/ml), 100 ng/ml recombinant human stem cell factor (rhSCF), 20 ng/ml recombinant human thrombopoietin (rhTPO), 100 ng/ml recombinant human Flt3 ligand (rhFlt3), and 20 ng/ml recombinant human IL6 (rhIL6) (all from Peprotech) 16 to 24 hours prior to transduction. HSPC were then transduced at a concentration of 1 × 10^6^ cells per milliliter with VSV-G-pseudotyped SINLV for 16 hours at the indicated multiplicity of infection (MOI) in the same medium. G-CSF mobilized peripheral blood CD34^+^ cells were maintained in culture in CellGro medium (Cell Genix) containing a cocktail of cytokines: 60 ng/ml IL-3, 100 ng/ml TPO, 300 ng/ml SCF, and 300 ng/ml FLT-3L (all from Cell Peprotech). Transductions were performed with the indicated dose of vectors for 16 hours in the same cytokine-containing medium. For single hit reporter LV transductions cells were washed and maintained in serum-free medium supplemented with cytokines as above until the reading of the percentage of positive cells by FACS. IMDM supplemented with 10% FBS, 25 ng/ml rhSCF, 5 ng/ml rhIL6-3, 25 ng/ml rhFlt3 and 5 ng/ml rhTPO was used for the seven days following the cytofluorimetry analysis before measurement of vector copy numbers. MDM were transduced 7-10 days after differentiation. To perform OE, KD and KO experiments cells were transduced with the respective LV and then challenged with a second transduction with or without drugs. Except for primary cells, KD and KO cells were selected with Puromycin before the second hit of transduction.

#### Compounds

Cyclosporin H, MG132, 3-MA, (all from Sigma-Aldrich), PP2 (Merk Millipore), Bafilomycin, GSK-LSD1 and Spautin-1 (Cayman Chemicals), were resuspended and stored following the manufacturer’s instructions. Any of these compounds were added to the transduction medium at the indicated concentration prior to transduction and washed out with the vector 16-20 hours later. Human IFNα (from pbl assay science 11105-1) pre-stimulation was performed for 24 hours together with human cytokines cocktail at the indicated concentration.

#### Flowcytometry

All cytometric analyses were performed using the FACSCanto III instrument and Cytoflex S instruments (BD Biosciences) and analysed with the FACS Express software (De Novo Software).

#### Transduced cells

GFP or BFP expression in transduced cells was measured 5 days post-transduction. Adherent MDM were detached using Trypsin-EDTA (Sigma), washed and resuspended in PBS containing 2% FBS. Cells grown in suspension were washed and resuspended in PBS containing 2% FBS. 7-aminoactinomycin D (Sigma Aldrich) were included during sample preparation according to the manufacturer’s instructions to identify dead cells.

#### Western blot

Whole cell extracts were prepared as previously described (67, 68).Samples were subjected to SDS-PAGE, transferred to PVDF membrane by electroblotting, and blotted with mouse polyclonal antibody CypA Ab (1:500 dilution, Santa-Cruz Biotechnology catalogue number sc-134310); mouse monoclonal antibody (Ab) raised against CypD/F (1:1000 dilution, Abcam catalogue number [E11AE12BD4] ab110324), rabbit anti-IFITM3 polyclonal Ab (1:1000 dilution, from proteintech catalogue number 11714-1-AP); mouse anti-Fyn monoclonal Ab (1:3000 dilution, from Novus Biologicals catalogue number NB500-517); rabbit anti-H3K9Me3 (1:1000 Abcam catalogue number ab8898); rabbit anti-LSD1 (1:4000 Diagenode catalogue number C15410067); An anti-Actin Ab (1:10000 Sigma-Aldrich catalogue number)was used as a normalizer.

#### Native PAGE

For native protein extraction, 3x10^5^cells were pelleted for each condition. Cells were lysed in Native Lysis Buffer (50mM Tris-HCl pH 7.5, 150 nMNaCl, 1mM EDTA, 1% NP-40) in presence of Protease cocktail inhibitor (cOmplete™, EDTA-free Protease Inhibitor Cocktail, Sigma Aldrich, catalogue number 11873580001). Protein lysates were left rotating on-wheel at 4 °C for 15 min and cleared by 15 min of centrifugation at 4°C at 13200 (max speed). Protein supernatants were transferred in new empty tube and quantified as previously described (Kajaste-Rudnitski et al, 2011; Kajaste-Rudnitski et al, 2006). For Native PAGE, 7% Running native gel and 4% Stacking native gel were prepared. Before running the native protein samples, the inner electrophoretic chamber was filled with fresh Native Running Buffer (25mM Tris, 192mM Glycin, pH 8.4) with 1% of Deoxycholate (DOC) and the outer chamber with Native Running Buffer without DOC and the gel was run empty at 4°C for 30 min at 40mA. 20ug of protein diluted in native sample buffer 3X (187,5mMTris-HCl, 45% Glycerol, 3% DOC and Bromophenol blue) were run at 4°C for 4 hours at 15 mA. Samples were transferred to a Nitrocellulose membrane by electroblotting and blotted with rabbit anti-IFITM3 polyclonal Ab (1:1000 dilution, from proteintech catalogue number 11714-1-AP), anti-Vinculin (1:20000 dilution Millipore) was used as a normalizer.

#### Immunofluorescence microscopy

5x10^4^ THP1, K562 or CBCD34^+^ cells were seeded in Multitest slide glass 10-well 8mm (from MP Biomedicals) pre-coated with poly-L-lysine solution (Sigma-Aldrich) for 20 minutes. Cells were fixed with 4% paraformaldehyde (in 1X PBS) for 20 minutes at room temperature, washed with 1X PBS and permeabilized with 0.1% Triton X-100 for 20 minutes at room temperature. For blocking non-specific sites cells were incubated 30 minutes in PBG (5% BSA, 2% gelatin from cold water fish skin, from Sigma-Aldrich) and then stained for 2 till 16 hours with rabbit anti-IFITM3 polyclonal antibody (1:200 dilution from proteintech catalogue number 11714-1-AP) and mouse anti-LAMP1 monoclonal antibody (1:100 dilution from Abcam [H4A3] (ab25630)). After 3 washes with 1X PBS, cells were incubated with donkey anti-Rabbit IgG, AlexaFluor 488 or 555 (1:500dilution from Thermo Fisher Scientific catalogue numberA-21206, A-31572) and anti-mouse IgG, AlexaFluor 647 (1:500 dilution from Thermo Fisher Scientific catalogue number A-31573), for 1-2 hours at room temperature. Nuclei were stained with DAPI (4′,6-diamidino-2-phenylindole; 10236276001, Roche, Basel, Switzerland) for 5 minutes at room temperature. Images were recorded using the TCS SP5 Leica confocal microscope, 60x with oil.

#### Bioinformatic analysis

Amino acid sequences of the human IFITM1, IFITM2 and IFITM3 proteins were retrieved from the RefSeq database (69) using the following protein accessions NP_003632.4, NP_006426.2 and NP_066362.2 respectively.IFITM3 protein sequences from other species were collected from orthologues of the human IFITM3 gene using gene annotations from the RefSeq database including representative species from mammals, birds and fish. The Refseq protein accessions were used to retrieve the amino acid sequences of the proteins from the different species analyzed: NP_001185686.1 (*Pan troglodytes*), NP_079654.1 (*Mus Musculus*), NP_001129596.1 (*Rattusnorvegicus*), NP_001188311.1 (*Susscrofa*), NP_001071609.1 (*Bostaurus*), NP_001336988.1 (*Gallus gallus*) and NP_001170783.1 (*Danio rerio*). Alignment of the amino acid sequences was performed using the Clustal Omega tool (version 1.2.4) with default settings (70). Protein domains were retrieved using the InterPro database (71). The multiple sequence alignment and the protein domains were then viewed and edited using Jalview (version 2) alignment software (72). Residues conservation was calculated as a numerical index reflecting the conservation of physico-chemical properties for each column of the alignment as previously described (73).Conservation is visualized on the alignment or a as a histogram giving the score for each column. Conserved columns have the highest score and are indicated by a ‘*’, and columns with mutations where all properties are conserved are marked with a ‘+’.

#### Statistical analysis

In all studies, values are expressed as mean ± standard error of the mean (SEM). Statistical analyses were performed by unpaired Mann Whitney test, as indicated. Percentages were converted into Log ODDs for statistical analysis. Differences were considered statistically significant at p<0.05.

## Acknowledgments

This work was supported by grants from the European Research Council (ERC-CoG 819815-ImmunoStem), The Italian Ministry of Health (GR-2018-12366006) and the Telethon Foundation (TELE20-C3) to AKR and by grants from the Medical Research Council (G1000196) and the Wellcome Trust (106223/Z/14/Z) to MHM. GU conducted this study as partial fulfilment of their Ph.D. in Molecular Medicine, Program in Cellular and Molecular Physiopathology, International Ph.D. School, Vita-Salute San Raffaele University, Milan, Italy. We wish to thank Andy Wang for technical help with plasmid cloning, Nathaniel Landau for reagents, Yves Gaudin for reagents and critical input and Cesare Covino from Alembic for the help with the confocal imaging acquisition and analysis.

## Authorship contribution

GU and AMSG conducted experiments and analyzed data. IC provided technical assistance. MAA and IM performed bioinformatics analysis. LA and MHM provided reagents and intellectual input. GU, CP and AKR designed the research studies, analyzed data, and wrote the manuscript.

## Declaration of Interests

Authors declare no conflict of interest.

## Supplementary Figure legends

**Figure S1. CsH degrades IFITM3 independently from autophagy. *A*-C.** THP1 over-expressing (Oe) IFITM3 or controls **(A-C)** and CB-HSPC **(B)** were transduced after an O/N treatment with CsH, the autophagy inhibitors 3MA and Spautin-1 or the combination of CsH with either one of the two compounds. Transduction was measured after five days by FACS. Data represent the mean ± SEM (n=3-5), p values are for Mann Whitney test. * for p<0.05. **B-D.** IFITM3 degradation by CsH was investigated by WB in THP1 after 16h treatment with CsH alone or in combination with one of the autophagy inhibitors. Representative images are shown. IFITM3 was quantified and normalized to Actin using ImageJ (n=2).

**Figure S2. IFITM3 single-lysine mutants localize in the endo-lysosomes.** Immunofluorescence images were performed using TCS SP5 Leica confocal microscope, 60x with oil on THP1 over-expressing IFITM3 or IFITM3 single lysine mutants after 16h treatment with CsH. IFITM3 intensity was quantified by ImageJ. Data represent the mean ± SEM of five experiments (n=12).

**Figure S3. CsH-mediated degradation of IFITM3 is independent from Lys88 methylation. A-B.** THP1 over-expressing IFITM3 or control **(A)** and CB-HSPC **(B)** were transduced with VSV-G LV at MOI 1 in presence or absence of CsH, LSD1 inhibitor or the combination of the two compounds. Transduction was evaluated by FACS at 5 days. Data represent the mean ± SEM of at least three experiments. **C.** LSD1 inhibition by the LSD1 specific inhibitor was investigated by WB in THP1 stimulated or not with IFNα after 16h treatment with CsH, LSD1 inhibitor or their combination (n=2). IFITM3 was quantified and normalized to Actin using ImageJ.

**Figure S4. Lysine-less IFITM3 mutant maintains CsH sensitivity.** IFITM3 or IFITM3 lysine-less degradation by CsH was investigated by IF using TCS SP5 Leica confocal microscope 60x with oil in THP1 and MDM after 16h treatment with CsH. Representative images are shown (n>12).

**Figure S5. The lysines in the intracellular loop of IFITM3 are not required for endolysosomal localization and CsH-mediated degradation. A.** IFITM3 wild-type or IFITM3 triple lysine mutant (in red) co-localization (purple areas marked by white arrows) with the lysosome associated membrane protein 1 (LAMP1) marker (in blue) was evaluated by immunofluorescence using TCS SP5 Leica confocal microscope 60x with oil in THP1 and MDM over-expressing cells. Zoomed images are shown for THP1. Representative images are shown (n=12). **B**. Degradation of IFITM3 wild-type and triple lysine mutant IFITM3 was measured after 16h treatment with CsH in THP1 and MDM by IF using TCS SP5 Leica confocal microscope 60x with oil. Representative images are shown (n>8).

